# Total lung capacity without plethysmography

**DOI:** 10.1101/395160

**Authors:** Ori Adam, Inon Cohen, Wai-Ki Yip, Robert J. Shiner, Peter Calverley, Zachi Peles, Adam LaPrad, Yoni Dagan, Robert Brown, Julian Solway, Jeffrey J. Fredberg

## Abstract

**Background:** Among the most basic measures of respiratory function is the total lung capacity (TLC). TLC is the pulmonary gas volume at maximal lung inflation, which is the sum of the volume of gas that can be exhaled –the vital capacity (VC)– and the volume of gas that cannot –the residual volume (RV). Determination of VC requires only spirometry whereas determination of RV or TLC requires body plethysmography, gas dilution or washout, or thoracic imaging, each of which is more complex than spirometry, and none of which is suited to routine office practice, population screening, or community medicine. To fill this gap, we describe here a new approach to determine TLC without plethysmography.

**Methods:** In a heterogeneous population of 434 volunteers (265 male, 169 female; 201 healthy, 170 with airflow obstruction, and 63 with ventilatory restriction), we determined TLC in the standard fashion using conventional body plethysmography (TLC_pleth_). In the same individuals, we also determined TLC in a novel fashion using the MiniBox ™ (TLC_MB_). To obtain TLC_MB_, population-based data from traditional spirometry together with flow-interruption transients were subjected to data mining and machine-learning to create for each individual subject an unbiased statistical determination of TLC.

**Results:** For the combined heterogeneous population, we found TLC_pleth_ = 1.02TLC_MB_ −0.091 L, adjusted *r*^*2*^=0.824. For the heterogeneous population as a whole, and for each subpopulation, TLC_MB_ closely tracked TLC_pleth_. For 26 healthy subjects measured on different days, the coefficient of variation for repeated measurements in was 3.3% for TLC_pleth_ versus 1.6% for TLC_MB_.

**Conclusions:** These results establish the validity and potential utility of a new method for rapid, accurate, and repeatable determination of TLC in a heterogeneous patient population, but without the need of a plethysmograph.

## INTRODUCTION

The most common test of pulmonary function is spirometry, in which the volume of air flowing into and out of the respiratory system is measured directly. Spirometry can quantify volume differentials such as tidal volume (V_T_), forced vital capacity (FVC), or expiratory reserve volume (ERV), but cannot measure absolute volumes such as residual volume (RV), functional residual capacity (FRC), or total lung capacity (TLC). Absolute thoracic gas volumes (TGVs), such as RV, FRC, and TLC are useful in the diagnosis and management of respiratory system diseases, but their measurement requires technologies that are more complex and labor intensive than spirometry. Accordingly, RV, FRC, and TLC are often not available in adult or pediatric office practices.

To measure absolute lung volumes, the ATS/ERS Consensus Statement identifies five methods: whole body plethysmography, multi-breath helium dilution, nitrogen wash-out, computed tomography, and chest radiography^1^ Among these, body plethysmography is used most commonly and is widely regarded as the gold standard.^1-8^ Since its inception by Dubois in 1956^9^, body plethysmography has remained simple in principle but inherently complex, capital intensive, and physically imposing in practice. The plethysmograph can be uncomfortable or intimidating for the patient enclosed within it and, moreover, is dependent upon a skilled technician for calibration, operation and maintenance. Gas dilution and gas washout are well-established alternatives to body plethysmography, but each presents its own technical challenges.

For these reasons, investigators have explored alternative avenues to determine absolute lung volumes by other means, but with no success. Respiratory system impedance, even when extended to a wide range of forcing frequencies, has been shown to be inadequate to infer absolute lung volumes in the individual subject^10-17^Similarly, forced expiratory maneuvers have been shown to be inadequate.^18^ These failures may be attributable in part to the fact that the dynamics of gas distribution within the human lung are complex, and especially so in obstructive lung disease. Moreover, data interpretation in these approaches often rests upon fitting data to idealized mathematical models wherein there exists a wide range of TGV values that might fit the data equally well. When this happens, no useful determination of TGV can be inferred and the problem of mathematical inference is said to be non-unique or ill-posed.

To determine TLC in the individual subject without using a plethysmograph, here we take a different approach. Across a heterogeneous population of volunteers we measured traditional spirometry together with flow-interruption transients using a MiniBox ™, described below. In the same individuals we also measured TLC using traditional body plethysmography (TLC_pleth_). To obtain an unbiased statistical determination of TLC from corresponding MiniBox ™ data (TLC_MB_) in the individual subject, we then used data mining and machine-learning. Because TLC_MB_ for the individual subject is based on population-based data mining and machine-learning rather than a direct physiological measurement, its main advantage is that it rests upon no idealizing assumptions concerning respiratory system structure or function. Nevertheless, in any given subject this approach was able to determine TLC in a fashion that is accurate and repeatable when compared to TLC_pleth_ but without the need of a plethysmograph.

## METHODS

The study comprised three parts. First, in a heterogeneous population of 300 qualified volunteers, as described below, we measured TLC in the conventional manner using body plethysmography. In these same volunteers, we measured conventional spirometry and flow-interruption transients using a desktop device called the MiniBox (PulmOne Advanced Medical Devices, Ltd., Ra’anana, Israel). Based upon these data, we used a statistical algorithm –the LASSO^19, 20^– to find the strongest set of statistical predictors of TLC_pleth_. We arrived at a final statistical model with which to calculate TLC from the statistical predictors, called TLC_MB_. Second, we validated this statistical prediction using N-fold cross-validation.^21, 22^ Third, to evaluate this statistical model still further, we compared TLC_MB_ against TLC_pleth_ in a prospective heterogeneous cohort of 134 qualified volunteers.

### Subject population

We recruited volunteers at 6 institutions (Soroka University Medical Center, Beer Sheva, Israel; Rambam University Medical Center, Haifa, Israel; Maccabi HaShalom, Tel-Aviv, Israel; Maccabi HaSharon, Kefar-Saba, Israel; Assaf HaRofeh Hospital, Tzrifin, Israel; Tel Aviv Medical Center, Tel-Aviv, Israel) under a research protocol approved by the Ethical Review Board of each. The prospective clinical study was registered with ClinicalTrials.gov (NCT 01952431).

The population comprised three groups (Table 1): 1) healthy subjects; 2) subjects with airflow obstruction, such as chronic obstructive lung disease (COPD) or asthma and with varying severity level (mild, moderate, and severe); and 3) subjects with restrictive ventilatory disorders. Patients were recruited from the Lung Function Laboratory at each institution. In each case, disease severity was defined by the criteria in ATS/ERS guidelines.^23^

**Table 1:** Basic Subject Characteristics.

| **Table 1: Basic Subject Characteristics.** |  |  |  |
| --- | --- | --- | --- |
| **Characteristic** | **Entire Dataset**<br>**(N = 434)** | **Initial Model Development**<br>**(N = 300)** | **Prospective Validation**<br>**(N = 134)** |
| Male | 265<br>(61.1%) | 185<br>(61.7%) | 80<br>(59.7%) |
| Female | 169<br>(38.9%) | 115<br>(38.3%) | 54<br>(40.3%) |
| Age (years)* | 45.8 ± 19.2 | 45.0 ± 20.1 | 47.8 ± 17.0 |
| Height (cm)* | 168.6 ± 9.9 | 168.2 ± 9.7 | 169.6 ± 10.4 |
| Weight (kg)* | 75.0 ± 17.3 | 73.6 ± 16.5 | 78.1 ± 18.8 |
| Plethysmographic TLC (L)* | 5.6 ± 1.4 | 5.6 ± 1.3 | 5.6 ± 1.5 |
| **Respiratory Condition: †** |  |  |  |
| Healthy | 201<br>(46.3%) | 150<br>(50.0%) | 51<br>(38.1%) |
| Obstructed | 170<br>(39.2%) | 114<br>(38.0%) | 56<br>(41.8%) |
| Restrictive | 63<br>(14.5%) | 36<br>(12.0%) | 27<br>(20.1%) |
| **Obstruction Severity: †** |  |  |  |
| Mild | 74<br>(17.1%) | 42<br>(14.0%) | 31<br>(23.2%) |
| Moderate | 240<br>(55.3%) | 179<br>(59.7%) | 62<br>(46.4%) |
| Severe | 120<br>(27.6%) | 79<br>(26.3%) | 41<br>(30.4%) |

\* Values are means ± SD.
† Defined according to ATS standards.<sup>22</sup>

Subjects were considered eligible if they: a) provided informed consent; b) were at least 18 years of age; and c) were cooperative and capable of following instructions. Healthy subjects were eligible if they: a) never smoked; b) had no known history of respiratory, cardiovascular, hepatic, renal or metabolic disease; c) had a BMI < 35 kg/m^2^; d) had no persistent (lasting greater than 3 days) respiratory symptoms during the last 12 months (e.g., dyspnea, chronic cough, wheezing or phlegm); and e) had no history suggesting upper respiratory infection during the three weeks prior to testing. Non-healthy subjects were eligible if they had a documented obstructive or restrictive respiratory disorder.

Subjects were excluded from the study if they: a) were pregnant at the time of the study; b) had performed any significant physical activity that has resulted in breathlessness during 1 hour prior to the study; c) had a tracheostomy; d) were unable to satisfactorily perform routine, full lung function testing including body plethysmography (e.g., due to claustrophobia or inability to perform the panting maneuver required for plethysmography); e) were unable or unwilling to give informed consent; or f) were unable to complete the protocol.

For each subject, all measurements were made in the same laboratory, by the same technician, and were competed within two hours. The technician also recorded the subject’s gender, date of birth, height, weight, and summary medical history.

### Body plethysmography

Depending on the study site, different commercial body plethysmographs were used: a) ZAN 500 (nSpire Health, Inc) - Soroka, Rambam, Maccabi HaSharon, Assaf HaRofeh; b) Platinum Elite-Series (MedGraphics) – Maccabi HaShalom; c) MasterScreen Lab (Erich Jaegar, CareFusion) – Tel Aviv, Assaf HaRofeh. Associated transducers were calibrated in accordance with the manufacturers’ user manuals. Device calibration and device agreement between institutions were verified using manually operated isothermal containers (3 L or 5 L) filled with copper wool, as well as by measuring a healthy control subject with a known TLC.

Body plethysmography measurements were performed in accordance with manufacturer recommendations and ATS/ERS guidelines. ^1^Subjects panted at FRC at 0.5 to 1 Hz against a closed valve and then inhaled to TLC followed by slow exhalation to RV. The thoracic gas volume (TGV) at FRC was calculated as the mean of the first 3 individual FRCs that were within 5% of each other in which the 2 highest inspiratory capacity (IC) measurements were within 10% (or 0.15 L) of each other. TLC_pleth_ was calculated by adding the largest of the three ICs to the mean TGV.

### MiniBox

The MiniBox™ (PulmOne Advanced Medical Devices, Ltd., Ra’anana, Israel) is a table-top unit that includes a spirometer and a flow-interruption device. The flow-interruption device (Figure 1A) consists of a rigid 16.3 L container, a rapidly closing valve (<10 msec), and a hotwire anemometer-type flowmeter (working range ± 5 L/s) (Figure 1B). Calibration of the MiniBox flowmeters was performed daily using a standard 3 L syringe.

**Figure 1:**
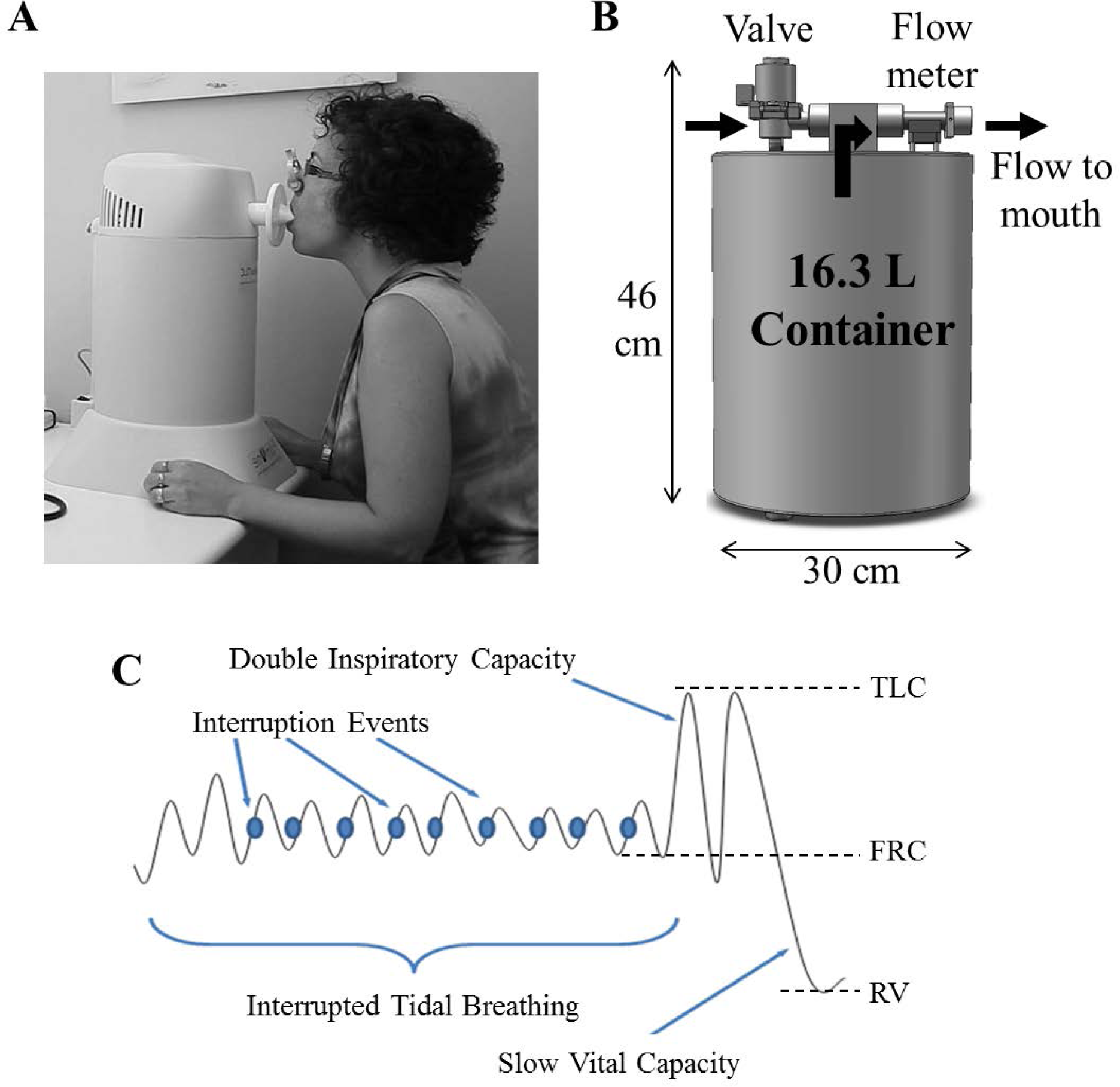
Overview of the MiniBoxPFT flow-interruption device and measurement procedure. **a)** Photograph of the MiniBox flow-interruption device during operation. The device sits on a table and the subject breathes tidally through a bacterial filter. **b)** Schematic illustration of the components of the MiniBox flow-interruption device, depicting the relative positions of the cylindrical container, valve, and flowmeter. Arrows indicate the direction of flow during inspiration. **c**) Schematic illustration of the breathing pattern required for lung volume measurement using the MiniBox flow-interruption device. Increasing lung volume is shown on the vertical axis and time increases to the right. During tidal breathing, brief interruptions are triggered in the vicinity of mid-inspiration (dots). After a minimum of 25 such interruptions or a maximum of 150 seconds, the subject then inhales maximally to TLC twice (double inspiratory capacity) and then exhales slowly to RV (slow expiratory vital capacity). In the illustration, only 9 interruptions are shown due to space limitations.

Subjects first performed spirometry using either the spirometer associated with the body plethysmograph or the hand-held spirometer associated with the MiniBox. For each participating laboratory, spirometry measurements were performed in accordance with the manufacturers’ user manuals and in a manner typical for that laboratory, as assessed by the local laboratory director. By design, spirometry measurements corresponded to real-world circumstances so as to include real-world methodological variability. That being the case, these measurements did not necessarily conform to ATS/ERS guidelines. ^24^ For example, in retrospective analysis we found that not all spirometry efforts continued for at least 6 seconds. At least 3 spirometry efforts were recorded. For SVC and forced vital capacity (FVC), additional maneuvers were performed until the 2 highest values of the 3 measurements comprising the selected group were within 5% (or 0.15 L) of each other. Similarly for IC, the 2 highest values of the 3 measurements comprising the selected group were required to be within 10% (or 0.15 L) of each other.

Subjects were next measured with the MiniBox flow-interruption device. With cheeks supported and a nose clip in place, each subject sat upright in a chair and breathed through a disposable bacterial filter attached to the MiniBox flow-interruption device (Figure 1A). The subject was asked to breathe normally for a short time until comfortable with the device. Then, brief flow interruptions (∼70 msec) were automatically triggered in the vicinity of mid-inspiration of each tidal breath (Figure 1C). After a minimum of 25 such interruptions or a maximum of 150 seconds of tidal breathing, the subject performed a maximal inspiration twice to reach total lung capacity (TLC). The subject then exhaled slowly to residual volume (RV).

The above flow-interruption measurement was repeated up to 3 times. The entire measurement was deemed acceptable if the SVC measured with the MiniBox flow-interruption device was within 10% (or 0.15 L) of the SVC measured with the spirometer. MiniBox flow interruption data were pre-processed and filtered based on pre-defined criteria.

### Statistical modelling

To construct an unbiased statistical model for TLC, and thus calculate TLC™ for each individual volunteer, we identified 137 plausible predictors of TLC from the spirometry and flow interruption data in 300 qualified volunteers. The metrics included conventional spirometry indices, transient flows and pressures measured at different time points during the flow-interruption, and their time derivatives. We then used a statistical algorithm – the LASSO^19, 25^ – to find the smallest possible set of predictors that produced a statistically significant determination of TLC. The LASSO is an extension of multiple linear regression that finds a combination of parameters while forcing all but a few coefficients to be precisely zero, thereby providing a minimal statistical model. Here, the LASSO was accomplished using a tunable parameter that constrains the coefficients with cross-validation using random sampling with replacement (bootstrapping)^26-29^ repeated 300 times for each value from a range of possible values. Each sampling was constrained according to the same ratio of male/female and healthy/non-healthy as the entire group of subjects. Using the LASSO applied to the dataset of 300 subjects, we arrived at a final statistical model to calculate TLC_MB_.

### N-fold cross-validation

To assess the predictive ability of the model, 10-fold cross-validation was used on the dataset of 300 subjects. ^21, 22^ The dataset was randomly divided using 10-fold for 50 times and the samples for each fold were selected randomly for each time. A 5-fold and leave-one-out cross validation (LOOCV) was also performed as comparison.

### Prospective validation

Last, we performed an independent prospective study to further validate the TLC_MB_ equation. In a prospective heterogeneous cohort of 134 additional volunteers not previously studied (Table 1), we repeated the protocol of MiniBox and body plethysmography measurements. We then used the new MiniBox data and the TLC_MB_ equation derived from the initial cohort of 300 subjects to calculate TLC_MB_ and compared it to TLC_pleth_ in the prospective cohort of 134 subjects.

## RESULTS

### Subject characteristics

A total of 564 subjects were enrolled, of whom 4 were unable to complete the protocol and 126 were disqualified based on quality assurance criteria. There were not any adverse events. The final qualified dataset comprised 300 subjects in the first cohort and 134 subjects in the prospective cohort (Table 1). Both cohorts included healthy individuals and patients with a range of diseases and a range of disease severities.

### TLC_MB_ versus TLC_pleth_

Across the entire mixed population of 300 qualified subjects, TLC_MB_ tracked TLC_pleth_ closely (Figure 2A; TLC_pleth_ = 1.02TLC_MB_ – 0.091 L, adjusted *r*^*2*^=0.824). In the subset of 150 healthy individuals, the variability was smallest (Figure 2B; TLC_pleth_ = 0.991TLC_MB_+ 0.0414 L, adjusted *r*^*2*^=0.852) while in the subset of 114 obstructed subjects (Figure 2C; TLC_pleth_ = 1.02TLC_MB_ – 0.004 L, adjusted *r*^*2*^=0.739) and in the subset of 36 restricted subjects (Figure 2D; TLC_pleth_ = 0.844TLC_MB_ – 0.474 L, adjusted *r*^*2*^=0.653), the variability was somewhat greater. Nonetheless, in each of these subpopulations, TLC_MB_ closely tracked TLC_pleth_.

**Figure 2:**
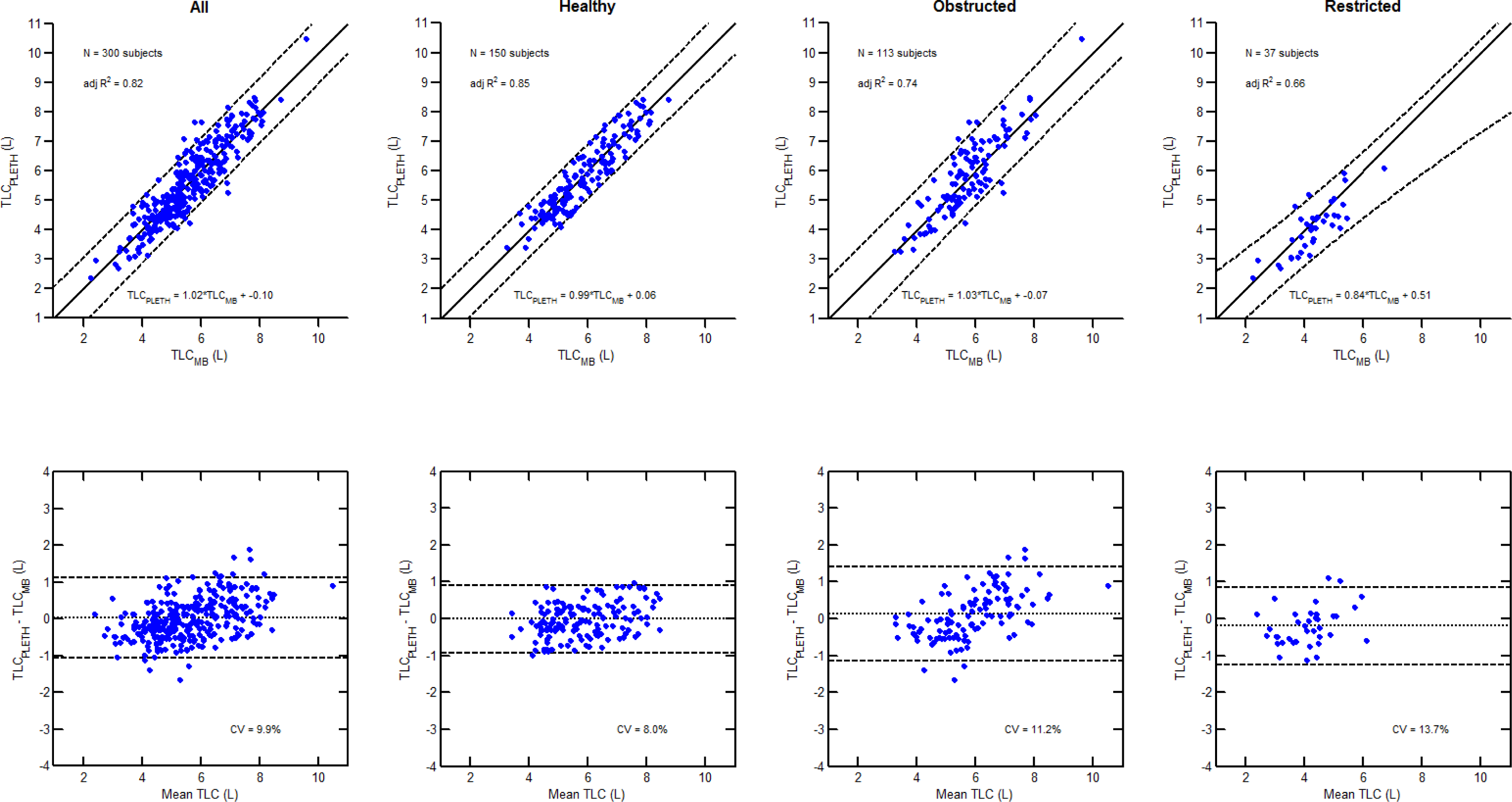
Comparison of TLC_MB_ and TLC_pleth_ in the initial 300 subject cohort. Top row: Scatter plots of plethysmographic TLC (TLC_pleth_) vs. MiniBox TLC (TLC_MB_) for all subjects **(A)**, healthy subjects only **(B)**, obstructed subjects only **(C)**, and restricted subjects only **(D)**. Males are represented by closed circles and females are represented by open circles. For subjects that were measured more than once on the device, the TLC is presented as the average value of all measurements. The dashed lines represent the unity line and the dotted lines represent the confidence intervals. The linear regression equation and the adjusted R^2^ are displayed within each graph. **Bottom row:** Associated Bland-Altman plots comparing TLC_MB_ to TLC_pleth_ for all subjects **(E)**, healthy subjects only **(F)**, obstructed subjects only **(G)**, and restricted subjects only **(H)**. The dotted lines represent the mean bias while the dashed lines represent the upper and lower limits (±1.96*SD). The coefficient of variation (CV) is displayed within each graph.

To examine differences between results from both methods and their dependence on lung size, we performed Bland-Altman analyses.^30^In the population as a whole (Figure 2E), and in each of the subpopulations (healthy - Figures 2F; obstructed - Figure 2G, and restrictive - Figure 2H), the coefficients of variations were 9.91%, 7.93%, 11.30%, and 13.70% respectively; the mean biases were small (0.01 L, −0.01 L, 0.11 L, and 0.20 L, respectively); also, there was no systematic trend of variability or bias with lung size.

### N-Fold cross validation

The mean prediction error using 10-fold cross-validation was 0.437 L with the mean prediction SE of 0.00171 L. The 5-fold and LOOCV also produced similar mean prediction errors. Thus, if the initial 300 volunteer cohort is representative of the population in which TLC_MB_ would be measured in practice, the statistical model has good predictive power.

### Independent validation in the prospective cohort

To further validate TLC_MB_, we then used the statistical model equation derived from the original cohort of 300 to calculate TLC_MB_ for each member of an independent prospective cohort of 134 (Figure 3). Although slopes and the adjusted *r*^*2*^ were slightly lower in that prospective cohort, TLC_MB_ closely tracked TLC_pleth_ and followed similar regression lines and confidence intervals.

**Figure 3:**
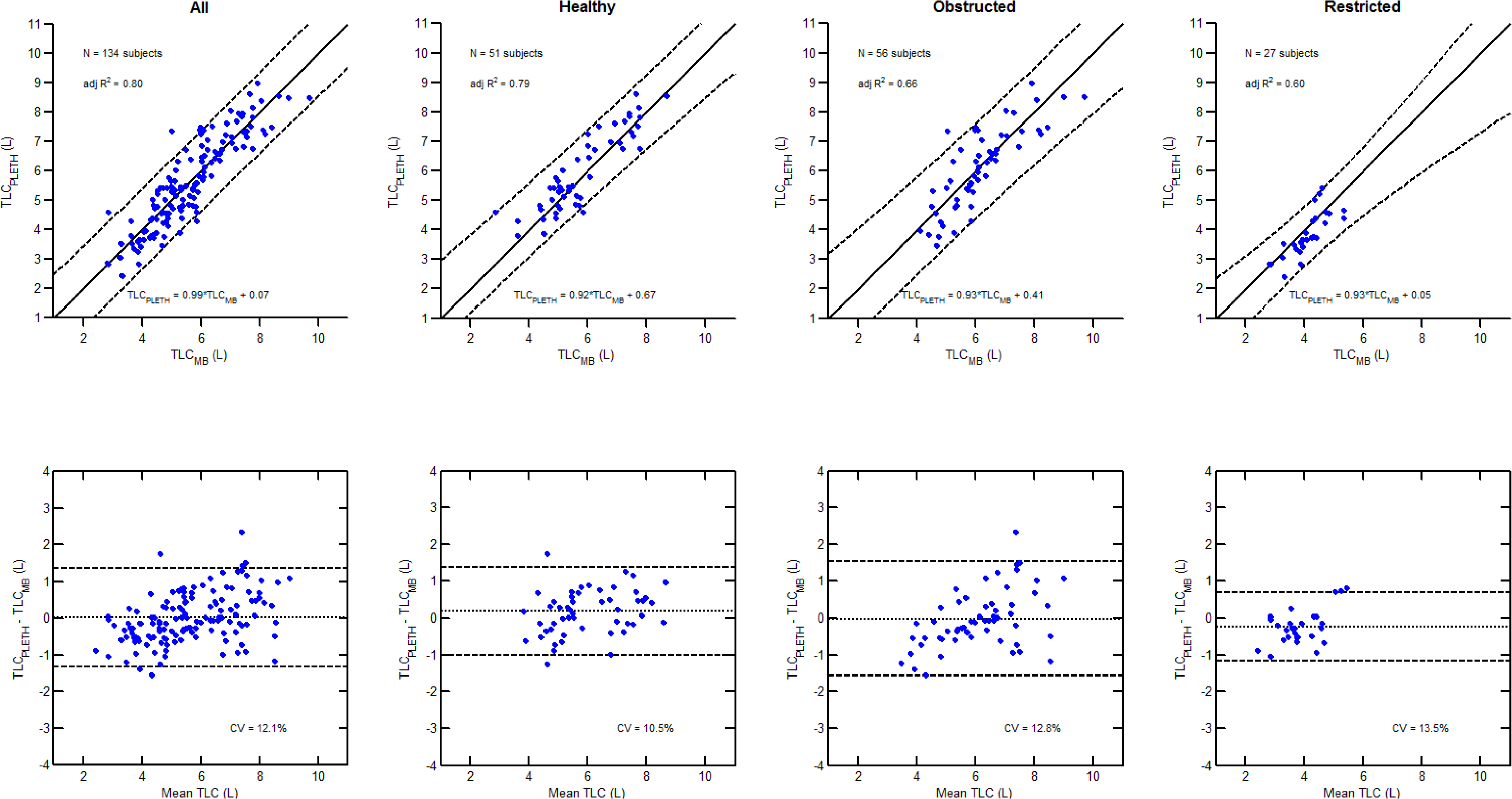
Prospective validation of TLC_MB_ in an independent cohort of 134 subjects. Top row: Scatter plots of plethysmographic TLC (TLC_pleth_) vs. MiniBox TLC (TLC_MB_) for all subjects **(A)**, healthy subjects only **(B)**, obstructed subjects only **(C)**, and restricted subjects only **(D)**. Males are represented by closed circles and females are represented by open circles. For subjects that were measured more than once on the device, the TLC is presented as the average value of all measurements. The dashed lines represent the unity line and the dotted lines represent the confidence intervals. The linear regression equation and the adjusted R^2^ are displayed within each graph. **Bottom row:** Associated Bland-Altman plots comparing MiniBoxPFT-derived TLC_stat_to plethysmographic TLC for all subjects **(E)**, healthy subjects only **(F)**, obstructed subjects only **(G)**, and restricted subjects only **(H)**. The dotted lines represent the mean bias while the dashed lines represent the upper and lower limits (±1.96*SD). The coefficient of variation (CV) is displayed within each graph.

Post hoc statistical analysis determined that the predictive contribution of both spirometry and flow interruption transients were statistically significant (p<0.01) and that spirometry contributed a majority of the predictive power.

### Day-to-day repeatability of TLC_MB_

From the initial 300 subject pool, we selected 26 healthy subjects at random to assess day-to-day repeatability with a minimum of 12 days between the measurements. Day-to-day repeatability was expressed as a coefficient of variation (CV; Figure 4). For TLC_MB_, the CV was 1.6% whereas for TLC_pleth_ the CV was 3.3%.

**Figure 4:**
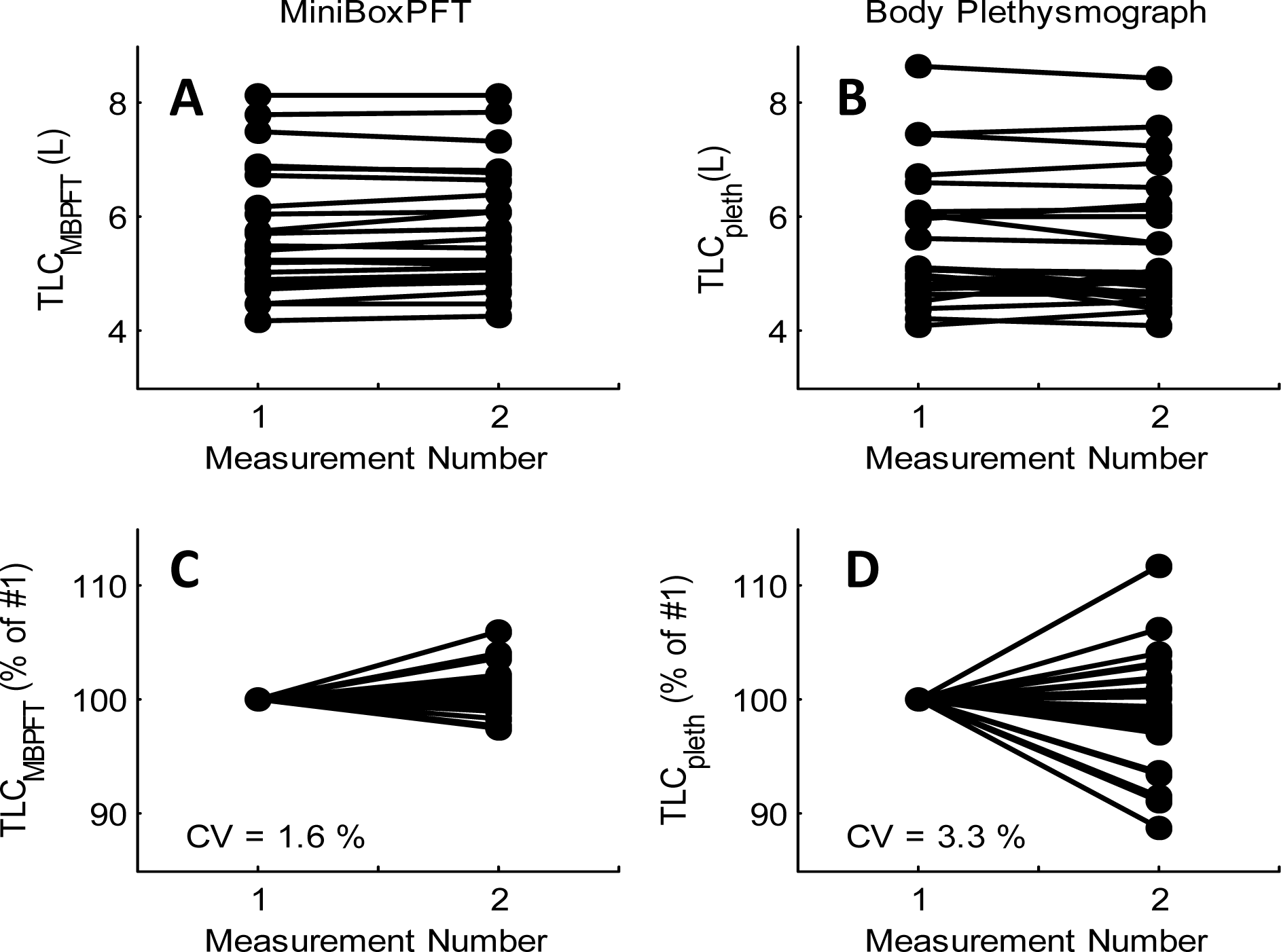
Repeatability of TLC determined using MiniBoxPFT or body plethysmography on two different days. Top row: TLC measured on day 1 and day 2 with the TLC_MB_ **(A)** and TLC_pleth_ **(B)**. **Bottom row (C, D):** TLC measured on day 2 normalized to the day 1 value. In 26 healthy subjects, the TLC_MB_ day-to-day repeatability was 1.6% compared to 3.3% for body plethysmograph.

## DISCUSSION

Based upon spirometry and flow interruption data taken together, here we apply data mining and machine learning to population-based observations in order to determine TLC in the individual subject. Within a highly heterogeneous population of subjects, we show that this approach yields accurate and reproducible determinations of TLC in the individual subject. Unlike the TLCpleth, the TLC_MB_ is not calculated based upon a physical principle or a mechanistic respiratory system model; physiological mechanisms were not a consideration. Instead, here we used inductive statistics and nonlinear systems identification, reminiscent of other applications of big data, to infer a relationship from which we could then make an accurate determination of TLC in the individual subject.

### Summary of clinical results

Across the entire population studied, across specific patient subgroups, and across a prospective heterogeneous population, our results show that TLC_MB_ is accurate compared to TLC_pleth_. Among our prospective cohort of 134 subjects, who were healthy or had varying severities of obstructive and restrictive diseases, TLC_MB_ correlated well with TLC_pleth_ (adjusted *r*^*2*^ = 0.795) with a slope close to unity (slope = 0.977) (Figure 3).

Furthermore, in a subset of healthy subjects, TLC_MB_ was appreciably more repeatable from day-to-day than was TLC_pleth_ (Figure 4), suggesting that TLC_MB_ might be particularly useful in longitudinal clinical management.

### Comparison to helium dilution and CT imaging

How does TLC_MB_ compare to other alternative technologies, such as gas dilution or computed tomography (CT), to measure absolute lung volumes? In a cohort of healthy, obstructive, and restrictive subjects, O’Donnell et al._^6^_ performed Bland-Altman analyses to compare TLC measured using both helium dilution (TLC_He_) and CT imaging (TLC_CT_) to TLC measured using plethysmography (TLC_pleth_).^6^ For TLC_He_ and TLC_CT_, the analysis showed coefficients of variation of 18.9% and 15.6%, respectively, together with systematic biases and trends for increasing error in subjects with larger TLCs (Figure 5B and 5C). Although we studied a different cohort, and results may therefore not be strictly comparable, Bland-Altman analysis of TLC_MB_ showed a coefficient of variation of 12.3% in our prospective cohort (N = 134 subjects), no systematic bias, and no trend of increasing error with increasing TLC (Figure 5A). While each of these technologies is based on a different mechanism-of-action, and thus is not expected to mimic plethysmographic TLC faithfully in all subjects, TLC_MB_ values had the smallest deviations from those of TLC_pleth_.

**Figure 5:**
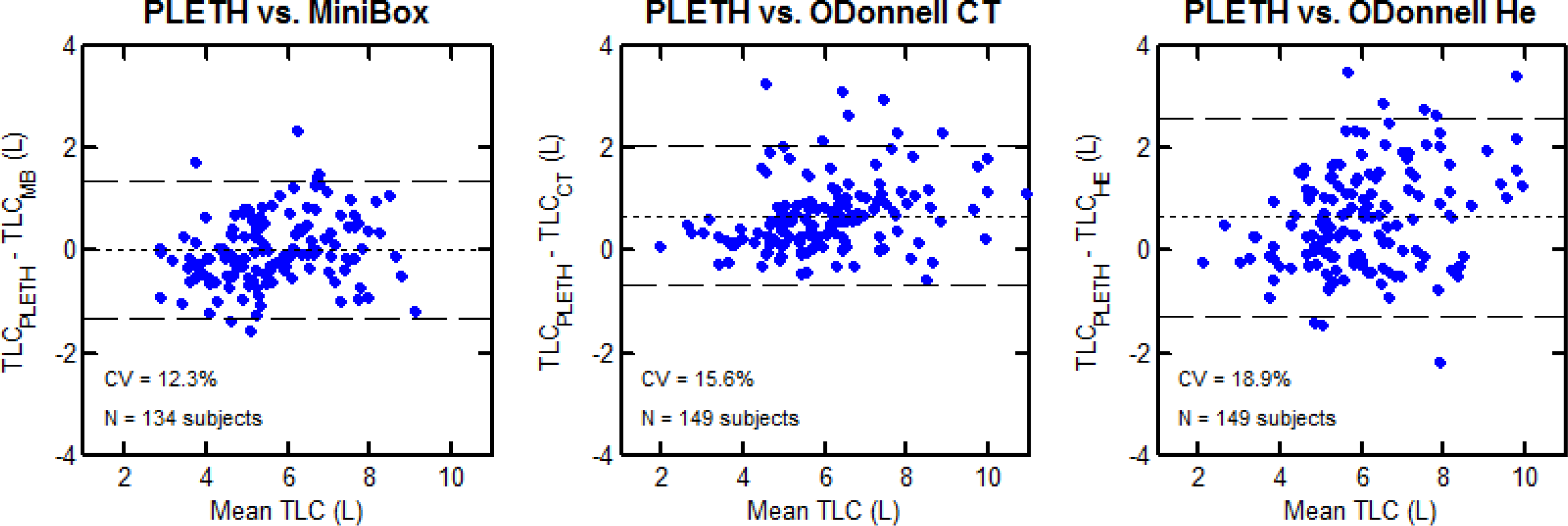
Bland-Altman Plots for TLC values determined by MiniBox, CT, or multibreath helium dilution, as compared to TLC by body plethysmography. In comparison to plethysmographic TLC, the absolute errors in MiniBoxPFT-derived TLC_MB_ are smaller than the errors in TLC determined using CT (TLC_CT_) or multibreath helium dilution (TLC_He_), as measured in O’Donnell et al. ^6^The dotted lines represent the mean bias while the dashed lines represent the upper and lower limits (±1.96*SD). The coefficient of variation (CV) is displayed within each graph.

### Limitations

The TLC_MB_ approach is limited in at least three important ways. First, TLC_MB_ is a population-based statistical approach. To the extent that specific populations might differ, different data training sets might be required. For example, pediatric populations, geriatric populations, or specific racial populations might require different training sets. Second, the flow interruption parameters improved the accuracy of TLCMB determinations by an amount that was small but nonetheless was highly significant statistically (p<0.01). Third, TLC_MB_ is designed to recapitulate as closely as possible TLC_pleth_. But TLC_pleth_ is itself subject to artifacts and is known to be an imperfect measure of TLC. ^31-34^ As such, any biases or errors inherent to TLC_pleth_ are necessarily inherent in TLC_MB_.

### Conclusions

The NHLBI, ATS, and ERS have encouraged innovation in technologies to measure absolute lung volumes so as to attain improved accuracy, ease of use, and rapidity of testing^35^, and have recommended rigorous testing to ensure no substantial differences in results compared with standard techniques. In all sub-populations tested, the MiniBox™ performed in a manner that compared favorably with body plethysmography. Also, the day-to-day variability of TLC_MB_ was smaller than that of TLCs derived from helium dilution, CT imaging, or body plethysmography. Accordingly, this study establishes the validity of TLC_MB_ for rapid, accurate, and repeatable determination of TLC in a heterogeneous population of healthy adults and those with respiratory system diseases.

## FOOTNOTES

### Previous Publications

Parts of this manuscript were presented previously in abstract form.

Author Contributions
O.A., R.J.S., P.C., R.B., J.S., and J.J.F. designed the study. R.J.S., P.C., R.B., J.S., and J.J.F. oversaw the study. O.A. and Z.P. performed the study. I.C., W.Y., and Y.D. analyzed the data. A.L., J.S., and J.J.F. contributed to the writing of the manuscript. All authors have seen and approved the final version of the manuscript.

